# Motivation to work for amino acids requires acute dopaminergic activity

**DOI:** 10.64898/2026.10.13.758662

**Authors:** Perham M Black, Kelcey T Stapleton, Geanette Lam, Aylin R Rodan, Adrian Rothenfluh

## Abstract

Aberrant motivation is a hallmark of many psychiatric disorders, and studying motivation in model systems can be useful to understand the neural mechanisms of these disorders. Having versatile, ethologically relevant assays with which to measure motivation is therefore important. However, motivational assays in *Drosophila* have not been able to measure willingness to exert effort to consume food. Here, we took advantage of the fact that flies prefer to feed on liquid over solid foods to develop a new assay using agarose as a barrier to feeding, which we termed the Fly Liquid vs Agarose Motivation Evaluation (FLAME). Unlike motivational assays based on overcoming bitterness, the FLAME measures willingness to work and is less confounded by deprivation acting on peripheral sensing of the modality to overcome. By depriving flies of specific nutrients and measuring their choice to overcome semi-solid agarose food to get those nutrients, we show that the FLAME is a versatile assay for measuring motivation. We show that flies choose to overcome agarose to consume sugar, salt, or amino acids, depending on the specific prior deprivation. We then show that show that flies rely on acute activity in a subset of their dopamine neurons during the choice period in order to work for amino acids when deprived. The FLAME thus allows the investigation of specific neural mechanisms of motivation.

**Significance Statement:** Having good assays to measure motivation in model systems is important for understanding psychiatric disease. In flies, existing assays have measured only some aspects of motivation, limiting progress. Here, we develop a better assay (the FLAME) for measuring motivation to feed in *Drosophila melanogaster*. We demonstrate the versatility of our assay and show that a subset of dopamine neurons in the fly are required for them to exert effort for amino acid consumption when deprived. The FLAME will therefore be of use to researchers seeking to understand the basic mechanisms of motivational drives.

## Introduction

Motivation is affected in numerous neuropsychiatric conditions where the drive for goals can either be excessively strong, as during drug addiction, or overly blunted, such as in depressive episodes. To delineate the neural mechanisms that regulate motivation, we need to be able to quantify motivation. One can think of motivation as a state where an animal is willing to pursue a goal, be that a nutrient, drug, or social interaction, at a cost. This cost can be represented by the animal’s willingness to overcome, or endure, overtly negative consequences in pursuit of the goal. Alternatively, the cost can be the exertion of effort and work for attaining the goal. In mammals, experimental costs used to quantify an animal’s motivation have included unpleasant or even painful stimuli, like bitter quinine or electric shock; alternatively, tasks are used in which animals need to exert physical effort by pressing a lever or by digging [Hoods 1961, Deroche-Gamonet 2004, Hopf 2010, Murdaugh 2024, Mendez-Vazquez 2026]. Having these complementary approaches allows for better experimental control to address confounds, such as reduced work effort due to motor fatigue, as opposed to lack of motivation, for example.

In *Drosophila*, overcoming bitterness as an aversive barrier to feeding has been the favored approach [Wu 2005, Devineni 2009, Sareen 2021]. While bitterness is an ethologically relevant feeding deterrent, its primary sensation at the periphery is downregulated when flies become hungry [Inagaki 2014, Le Due 2016, Mishra 2021]. This makes sense, ecologically, as hungry beggars can’t be picky choosers. However, this depotentiation of bitterness distorts the measure of motivation, as in hungry flies bitterness is ‘overcome’ both by motivation to endure the aversive cost of bitterness, but also by reduced primary perception of that very cost. To address this, we developed the FLEA [Mishra 2021], which uses electric shock as an aversive barrier to feeding that is not depotentiated by hunger. Electric shock has also been used as a cost to determine flies’ motivation for choosing a learned preferred odor [Jovanoski 2023]. Measuring work effort in *Drosophila* is much less common but includes the assessment of running speed on a ball in pursuit of an odor, which is a type of effort exertion [Sayin 2019], although this assay lacks a consummatory endpoint, however, and is therefore prone to extinction of the motivated behavior, which also confounds the quantification of motivation [Todd 2014].

To establish an assay to quantify *Drosophila’s* willingness to work, we developed the Fly Liquid vs Agarose Motivation Evaluation, or FLAME. In this two-choice dye feeding assay flies choose between interspersed liquid and semi-solid agarose feeding options, capitalizing on the fact that *Drosophila melanogaster* generally prefer to feed on softer foods [Jeong 2016, Zhang 2016, Sanchez-Alcaniz 2017]. Liquid food allows flies to continuously drink while keeping their proboscis in the liquid [Mishra 2021], whereas on solid food they repeatedly probe and suck, as they lack the equipment for biting off and masticating [Itskov 2014, Edgecomb 1994]. We replicate the previously reported preference for softer foods in the FLAME and then go on to show that flies increasingly consume semi-solid agarose options when offered extra external incentives in the agarose and when internal states change the value of such extra incentives.

We show that different nutrients can motivate semi-solid overcoming, and that the motivated state is nutrient-specific. Lastly, we show that acute dopaminergic neuronal activity is required to motivate amino acid consumption. The FLAME is therefore a versatile assay to investigate the neural mechanisms of consummatory motivation.

## Materials and Methods

### Fly Keeping

Flies were raised on standard food supplied by Archon Scientific, Durham, NC. (Archon Corn Syrup Soy after Nov 2024, Glucose before Nov 2024). Wild-type flies (*w* Berlin*) were raised at 25°C and 50% humidity in a 12h light cycle or at room temperature with ambient humidity and light. Wild-type flies were between 2-7 days old for all behavioral assays. *TH-GAL4>shibire^ts^* flies and their *+>shibire^ts^*controls were created by crossing *w*;TH-GAL4* flies to *w* Berlin* flies and then collecting their virgin offspring to cross to *w*;UAS-shibire^ts^*males to create sibling controls of equal genetic background that were sorted by eye color 1-4 days post eclosion before deprivation. These flies were kept at 20°C in a 12h L:D light cycle. The same protocol was followed for *PAM-GAL4* (*R58E02*).

### Two Choice Assay

Two-choice fluorescent dye feeding assay was adapted from a previously published assay [Peru y Colón de Portugal, 2014] using 60 well plates. In all two-choice experiments, flies were allowed to feed from either yellow (0.06% Fluorescein sodium salt, Sigma-Aldrich, St. Louis, MO) or pink-colored (0.01% Rhodamine B, Thermo Fisher Scientific, Waltham, MA) food options, with one color in the agarose option and the other in the liquid option. The color in agarose was counterbalanced in different plates to avoid any potential color-preference confound. A “50/50 Plate” was run in parallel, where only one (liquid) choice was available with both colors mixed in at the same time, to normalize fluorescent readings for preference indexes of 0, i.e. no preference. Upon 18 or 6 h wet starvation, flies were given 4ml of water on a fresh flug (vial enclosure) between 30 and 45 minutes before assay time to ensure dehydration was not a confound for all behavioral assays. Flies in the FLAME fed for 30 minutes in the dark at ambient temperature (21-23C) and humidity. For thermogenetic experiments, the plates were placed in an incubator set to the indicated assay temperature at 50% humidity. At the conclusion of the 30 min assay period plates were placed in a −80°C freezer, and then later (up to 72h) flies of a specific sex were placed in groups of three (for an n=1) into wells of a 384-well plate (Greiner Bio-One, Monroe, NC) to be scanned in the plate reader (BioTek Syngergy HTX, Agilent, Santa Clara, CA). Plates were excited at 485nm and emission was read at 528nm for yellow dye, and excited at 530nm and emission read at 590nm for pink dye.

Raw fluorescence data points were converted to preference-indexes by using the following method: 3% of the pink 530nm signal was subtracted from the 485nm yellow signal to correct for signal bleed-through. Then, the ratio of pink to yellow signal was determined in the 50/50 plate and used to normalize the colors. After normalization, a standard preference index was calculated: A-L/A+L, where A was the reading in the agarose wells, and L in the liquid wells. R code for data analysis is available upon request. Synergy HTX protocol is available upon request.

### Deprivation Conditions

Wet starvation: 25-50 flies were starved in narrow vials with 4ml of 0.7% Agar. Protein deprivation: 25-50 Flies (half male half female except for experiments with virgin females) were deprived in narrow vials filled with 4ml of 100mM Sucrose in 0.5% agarose and compared with protein-satiation vials filled with 4ml of 0.5% agarose, 100mM sucrose, and 1mM casamino acid mix solution. Salt deprivation: 25-50 flies (half male half female except for experiments with virgin females) were deprived in narrow vials with 4ml of 0.5% agarose and 100mM Sucrose solution, compared with 4ml of 0.5% agarose, 100mM sucrose, and 50mM NaCl for non-deprived flies. Protein deprivation with plenty of salt: Approximately 50 flies (half male half female) were deprived in narrow vials filled with 4ml of 0.5% agarose, 100mM Sucrose, and 50mM NaCl, and compared to flies in narrow vials with 100mM Sucrose, 1mM casamino mix, 0.5% agarose, and 50mM NaCl. Protein deprivation alongside 3iY treatment: Approximately 50 flies were deprived in narrow vials filled with 4ml of 0.5% agarose, 100mM Sucrose solution, 1mg/ml 3iY, and 0.5% acetic acid (solvent for 3iY) and compared with protein-satiation vials filled with 4ml of 0.5% agarose, 100mM sucrose, 1mM casamino acid mix, 1mg/ml 3iY, and 0.5% acetic acid. 3iY stands for 3-iodo-tyrosine and is an inhibitor of tyrosine hydroxylase.

### Experimental Design and Statistical Analyses

Preference index data is constrained at both −1 and 1 which renders the majority of data non-normally distributed, therefore non-parametric analyses were carried out. The Mann-Whitney U Test was used throughout for comparisons between two groups, and the Kruskal-Wallis test and Dunn’s multiple comparison test were used when comparing three groups. For interaction-design experiments, we used 2-way ANOVA with Sidak’s post-hoc comparison. No prior formal statistical analysis was employed to determine group size, however an n of between 40 and 64 is standard in our lab for fluorometric preference data. All groups of flies shown on the same graph were raised in the same conditions and experiments were run in parallel, or on the same day in the case of 6 vs 18hr starvation experiments. All experiments were repeated on at least two days and data were pooled.

## Results

### Agarose is a non-discounted effort-based barrier to feeding that can be overcome with external incentives and altered internal state

One way to measure motivation is by asking whether flies are willing to overcome bitterness to consume palatable food [Wu 2005, Devineni 2009, Sareen 2021]. However, the peripheral perception of bitterness is downregulated with hunger, [Inagaki 2014, Le Due 2016, Mishra 2021] introducing a confound to such assays. We thus asked whether the perception of agarose – as a food barrier of hardness to overcome – would be discounted by hunger. We observed no difference in avoidance of 1% agarose (*p*=0.99, 2-way ANOVA, Sidak’s post-hoc comparison) between males starved for 6 or 18h, and the same was true for 2% agarose (*p*=0.51; Fig.1A).

**Figure 1.**
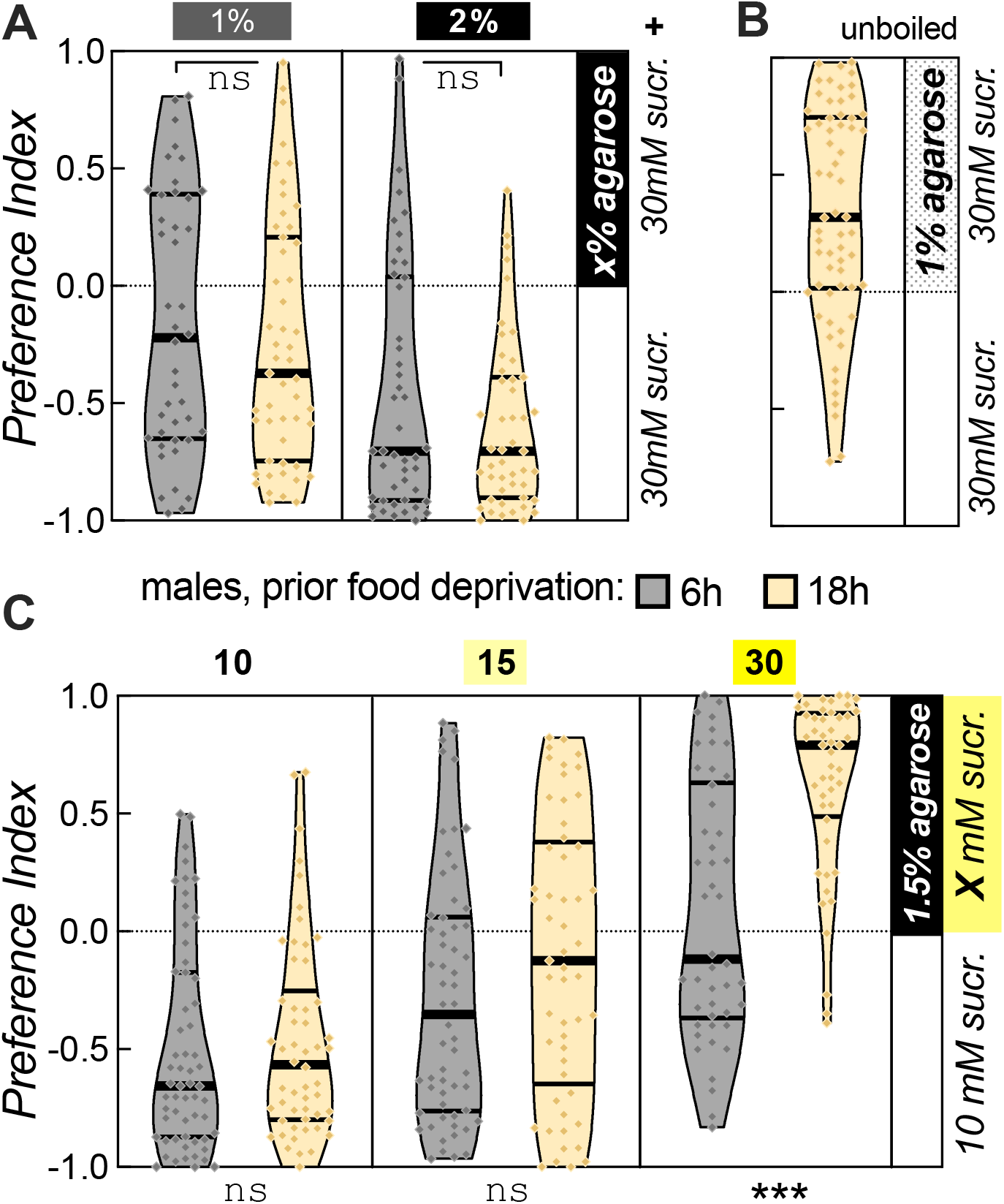
Starvation and extra sucrose lead to willingness to exert feeding effort. **A** Preference for 30mM sucrose in liquid vs in 1% agarose is no different in 18h (yellow) or 6h-food deprived (grey) male wild-type flies (*w* Berlin* for this and subsequent experiments; left). The same is true for 2% agarose (right). Groups: 6h starved 1% agarose (n=42), 18h starved 1% agarose (n=51); 6h starved 2% agarose (n=46), 18h starved 2% agarose (n=47). **B** Non-negative preference for 18h starved male wild-type flies for 30mM sucrose solution containing 1% unboiled agarose vs 30mM sucrose solution without agarose (n=62). **C** Interaction between food deprivation and sucrose incentive in the choice between 10mM liquid sucrose and different sucrose concentrations in hard agar. Only 30mM sucrose drove preference of hard sucrose consumption in 18 h but not 6 h-deprived males. Groups: 10mM 6h (n=59), 10mM 18h (n=54); 15mM 6h (n=58), 15mM 18h (n=53); 30mM 6h (n=46), 30mM 18h (n=48). (For this and all subsequent plots, each dot represents three flies that were scanned together during fluorescence reading. All data points for each experiment are shown along with median and quartiles on violin plots. Significance is displayed as \*\*\**p*≤0.001, \*\**p*<0.01, \**p*<0.05, ns *p*≥0.05, with exact p values and detailed specific test statistics available in the body of the paper.)

There was, however, a significant overall effect of agarose concentration (*F*_(1,159)_=19.5, *p*<0.001), suggesting that agarose avoidance is parametric, and the larger its concentration, the stronger the avoidance. To ascertain that it was indeed the hardness of agarose that was avoided, we also assayed the avoidance of 1% unboiled, and thus unhardened agarose and found that males actually preferred it over liquid sucrose without agarose (*p*<0.001, one-sample Wilcoxon test), suggesting that is the hardness of agarose that serves as a deterrent for consumption. Similar results were obtained for female flies (Figure S1A,B).

We next wanted to know whether an external incentive, such as additional sucrose, would act as an incentive for flies to overcome agarose. We thus pitted 10mM sucrose against 10, 15 and 30mM sucrose in 1.5% agarose (Figure 1C), and found a very strong main effect of sucrose (2-way ANOVA, *F*_(2,312)_=79.2, *p*<0.001), as well as of food-deprivation duration (*F*_(1,312)_=22.0, *p*<0.001). Interestingly, there was also a strong interaction between these two factors (*F*_(2,312)_=7.7, *p*<0.001), and post-hoc comparisons revealed that hungry males were not incentivized to overcome agarose for 15 vs 10mM sucrose (*p*=0.18, Sidak’s multiple comparison test), while they strongly preferred 30mM hard over 10mM liquid sucrose (*p*<0.001). Thus, external incentives can motivate flies to overcome food hardness, and internal states such as hunger can strongly potentiate this drive. Similar results were obtained for females (Figure S1C).

### Salt-deprivation and mating drive flies to overcome agarose to obtain NaCl

We next wanted to know whether our assay could also measure the drive to overcome agarose for nutrients other than sugar. Initial experiments suggested that NaCl was not as strong of an incentive as sugar (data not shown), and we therefore used a lower concentration of 0.7% agarose as a barrier to overcome in these experiments. We first asked whether 5 days of salt deprivation (see Methods) would change 0.7% agarose avoidance, and it did not in males (*p*=0.17, Mann-Whitney U test; Fig.2A), whereas in females it led to a significant (*p*=0.013) but small decrease (0.09 absolute difference in median avoidance) decrease in agarose avoidance (Fig.2C). We then incentivized the agarose side with 25 mM NaCl and that made deprived males overcome agarose to indifference, significantly different from agarose-avoiding undeprived males (*p*<0.001, Mann-Whitney U test; Fig.2B). We then performed the same experiment with females, but because females have been shown to change their salt intake with mating status [walker 2015] we assayed both mated and virgin females (Fig.2D) and found a main effect of mating status (*F*_(1,255)_=7.2, *p*=0.008) and a strong main effect of salt deprivation (*F*_(1,255)_=27.9, *p*<0.001). Both virgin (*p*=0.009, Sidak’s multiple comparison test) and mated females (*p*<0.001) were significantly less deterred by agarose to consume 25mM NaCl after 5d of salt deprivation, showing that different incentives, as well as distinct types of nutrient deprivation along with mating status can drive flies to overcome hard food in the FLAME.

**Figure 2.**
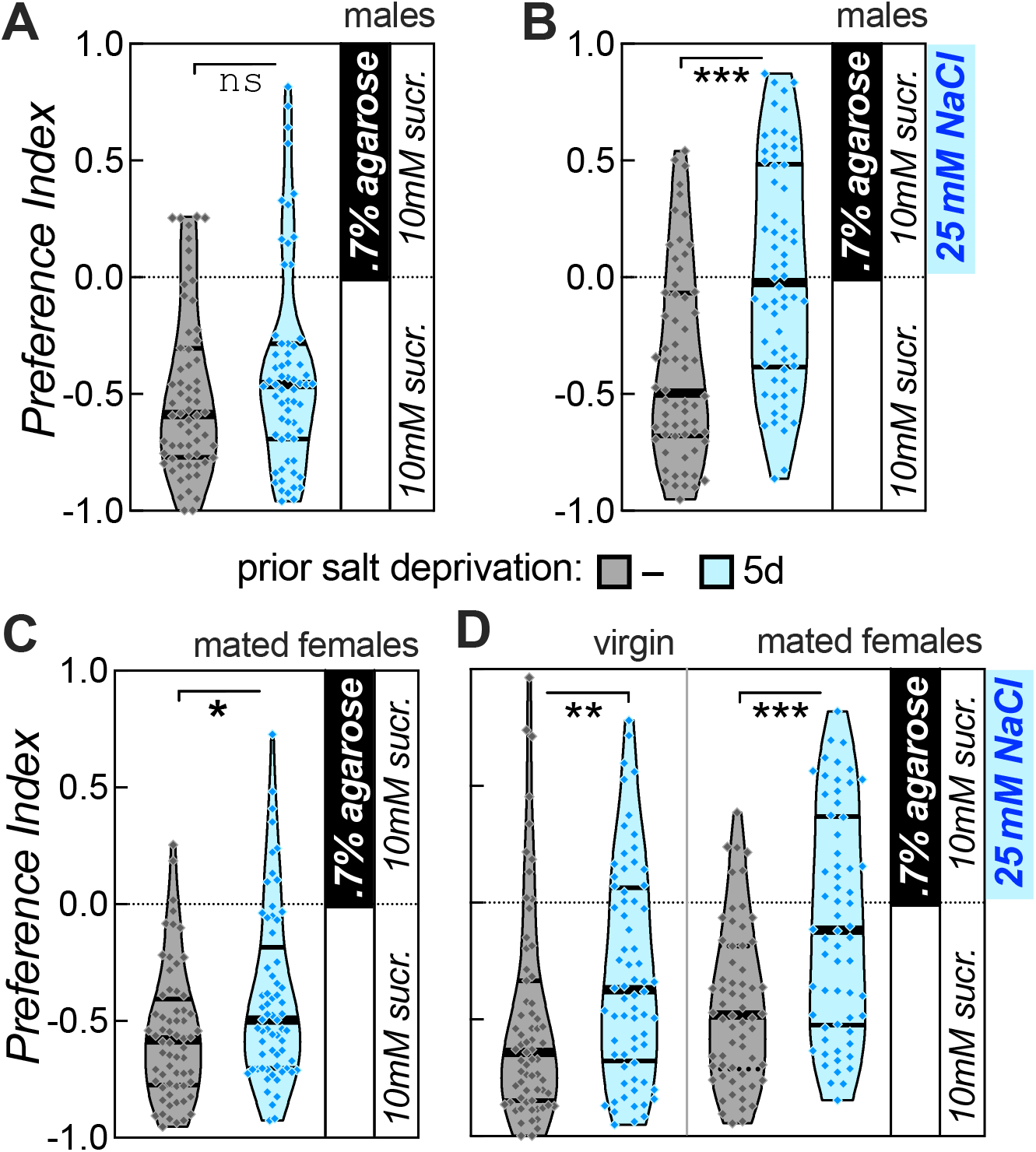
Salt deprivation drives flies to overcome agarose for salt consumption. **A** Avoidance of 0.7% agarose in male wild-type flies with 5-day salt deprivation does not change compared to those with no prior deprivation. Groups: no prior deprivation (n=64), salt deprived (n=63). **B** Preference increases for 0.7% agarose with a 25mm NaCl incentive in male wild-type flies with 5-day salt deprivation compared to those with no prior deprivation. Groups: no prior deprivation (n=64), salt deprived (n=66). **C** Avoidance of 0.7% agarose in female wild-type flies with 5-day salt deprivation decreases slightly, but significantly compared to those to no prior deprivation. Groups: no prior deprivation (n=64), salt deprived (n=65). **D** Preference increases for 0.7% agarose with a 25mM NaCl incentive in both virgin (left) and mated (right) female wild-type flies with 5-day salt deprivation compared to those with no prior deprivation. Groups: virgin no prior deprivation (n=67), virgin salt deprived (n=68); mated no prior deprivation (n=62), mated salt deprived (n=62).

### Protein Drive is synergistically regulated by mating status and deprivation

Unlike NaCl, amino acids (aa) proved to be highly motivating in our preliminary experiments (data not shown), and we therefore used very hard 3% agarose as the barrier in experiments using aa as an incentive. 3d of aa deprivation had no effect on agarose avoidance in males (*p*=0.13, Mann-Whitney U test; Fig.3A) or females (*p*=0.76; Fig.3C). We then aa deprived flies for 3d and incentivized the hard side only with 2mM aa (Fig.3B), this drove deprived males to indifference, significantly avoiding aa-containing agarose less than undeprived males (*p*<0.001). Like NaCl, female aa drive is known to depend on mating status [Vargas 2010, Ribeiro 2010], and while there was no main effect of mating status (*F*_(1,215)_=2.4, *p*=0.13), there was a strong effect of deprivation (*F*_(1,215)_=44.5, *p*<0.001) and a significant interaction (*F*_(1,215)_=10.9, *p*=0.0011) that drove mated females to prefer the hard side with 2mM aa (*p*<0.001, Sidak’s multiple comparison test), while virgin females significantly avoided hard 2mM aa less when deprived (p=0.04), but did not prefer the hard 2mM aa side (Fig.3D). These data show that aa deprivation and aa incentive strongly impact feeding drive and can motivate females to overcome hard agar (Fig.3D), which is normally a strong deterrent to feeding (Fig.3C).

**Figure 3.**
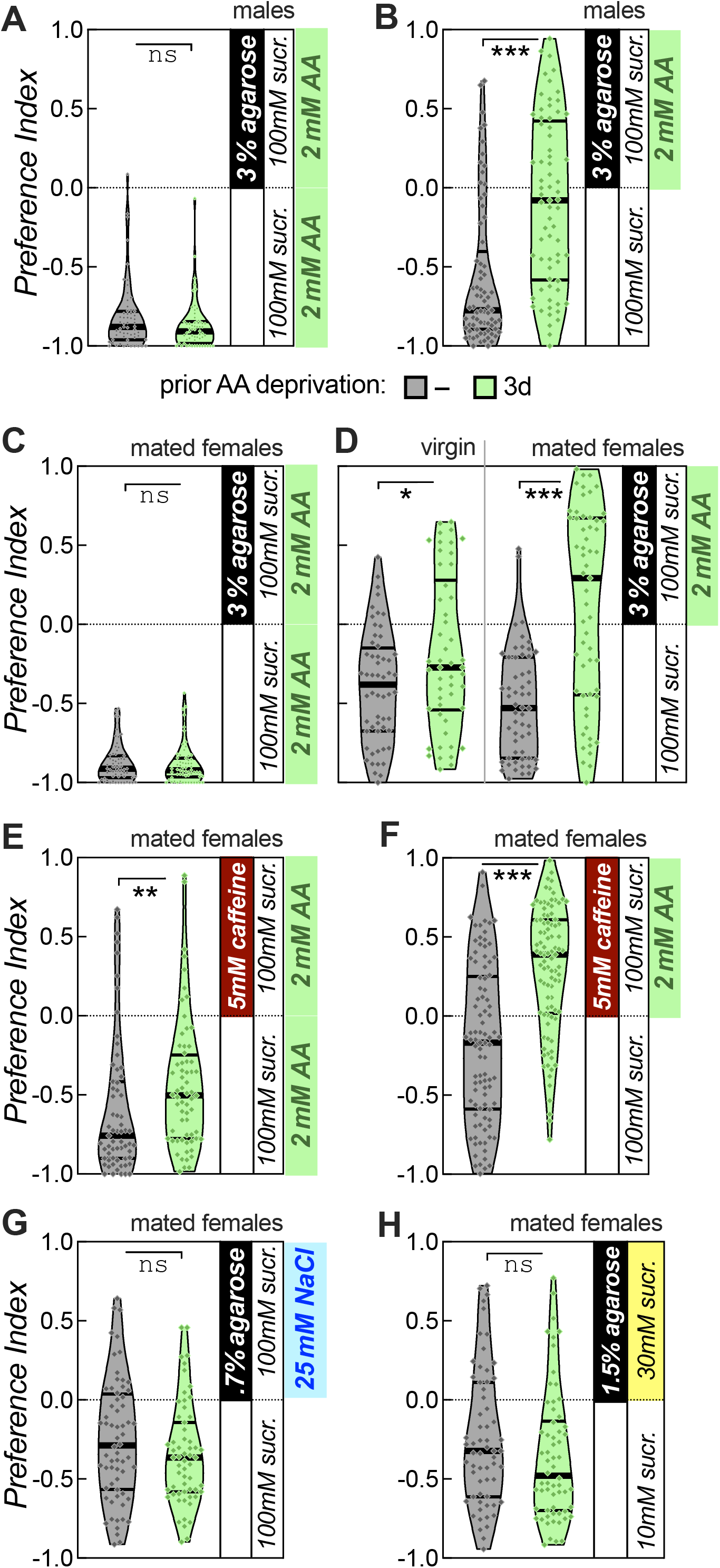
Protein deprivation selectively regulates amino acid motivation. **A** Avoidance of 3% agarose does not change in male wild-type flies with 3-day protein deprivation compared to those with no deprivation. Groups: no prior deprivation (n=59), protein deprived (n=58). **B** Preference increases for 3% agarose with 2mM amino acid incentive for male wild-type flies with 3-day protein deprivation compared to those without prior deprivation. Groups: no prior deprivation (n= 59), protein deprived (n=54). **C** Avoidance of 3% agarose does not change in mated female wild-type flies with 3-day protein deprivation compared to those with no deprivation. Groups: no prior deprivation (n=63), protein deprived (n=62). **D** Preference increases for 3% agarose with 2mM amino acid incentive for virgin and mated female wild-type flies with 3-day protein deprivation compared to those without prior protein deprivation. Groups: virgin no prior deprivation (n= 56), virgin deprived (n=50); mated no prior deprivation (n=56), mated deprived (n=57). **E** Caffeine avoidance is reduced by protein deprivation for mated female wild-type flies. Groups: no prior deprivation (n=63), protein deprived (n=64). **F** Caffeine is overcome by protein deprived female wild-type flies in an assay with 2mM aa incentive. Groups: no prior deprivation (n=93), protein deprived (n=94). **G,H** 3-day protein deprivation does not motivate flies to overcome agarose to consume extra salt, **G**, or extra sucrose, **H**. Groups in salt experiment: no prior deprivation (n=64), protein deprived (n=59); in sucrose experiment: no prior deprivation (n=62), protein deprived (n=62).

Previous investigations have used bitter taste as a barrier to overcome in order to assay motivation. We therefore wanted to compare the FLAME with an assay where flies had to overcome 5mM caffeine instead of 3% agarose. First, we asked whether aa deprivation would impact caffeine avoidance in the absence of extra aa incentive mixed with caffeine. Unlike unchanged agarose avoidance, aa deprivation significantly reduced mated females’ caffeine avoidance (*p*=0.003, Mann-Whitney U test; Fig.3E), suggesting that just like wet starvation [Inagaki 2014, Le Due 2016, Mishra 2021], deprivation of aa alone impacts the perception and/or processing of bitterness. Nevertheless, we then assayed aa deprivation as a driver to overcome caffeine for aa consumption, and it led to strong preference of bitter aa, compared to indifferent undeprived females (*p*<0.001; Fig.3F). Still, the effect of the 2mM aa incentive in caffeine is harder to quantify than when in agarose, given that caffeine avoidance is affected by the aa deprivation.

To test whether aa deprivation would affect drive for incentives other than aa, we aa deprived mated females in the presence of 50mM NaCl and then asked whether that would drive them to overcome agarose for 25mM salt. Deprivation of aa had no significant impact (*p*=0.21, Mann-Whitney U test; Fig.3E), and aa deprivation in the presence of 100mM sucrose similarly had no significant effect on preference for hard 30mM sucrose over liquid 10mM sucrose (*p*=0.11; Fig.3F). We obtained similar results with male flies (data not shown). Therefore, aa deprivation specifically drives flies to overcome agarose for aa, but not for other external incentives.

### Acute dopamine activity is required to drive flies to work for amino acids

Because aa drive in mated females was the strongest of the three drives assayed, we put our focus there and next asked whether dopamine is involved. We fed flies 3iY, which inhibits the rate-limiting dopamine synthesis enzyme tyrosine hydroxylase (TH), during the 3d aa deprivation, and then assayed them for drive to feed from hard aa (Fig.4A). Deprivation was a significant main effect (*F*_(1,179)_=15.2, *p*<0.001), and while the treatment was not a main effect (*F*_(1,179)_=0.001, *p*=0.97), there was a borderline significant interaction between treatment and deprivation (F_(1,179)_=3.7, *p*=0.055). Furthermore, deprivation incentivized control females to avoid hard aa less (*p*<0.001, Sidak’s multiple comparison test), but not 3iY-treated females (*p*=0.31), suggesting that dopamine is involved in aa drive.

**Figure 4.**
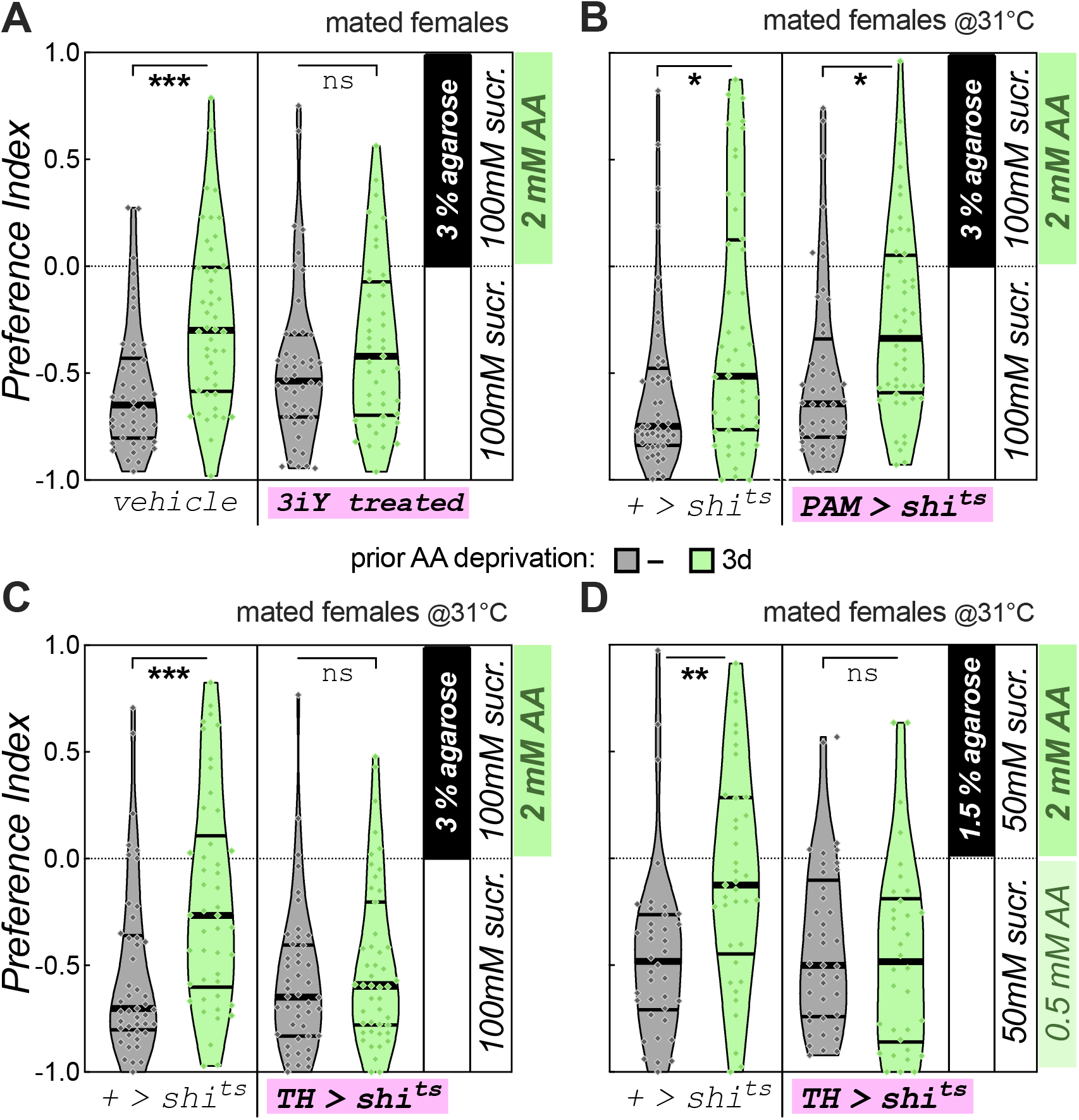
Dopaminergic requirement for amino acid motivation. **A** 3iY administration blocks deprivation-induced increase in preference for 3% agarose with 2mM aa incentive in mated females. 3iY was administered during the 3d-deprivation period. Groups: no prior deprivation given vehicle (n=46), protein deprived given vehicle (n=46); no prior deprivation given 3iY (n=46), protein deprived given 3iY (n=45). **B** Acute silencing of PAM dopamine neurons (*R58E02-Gal4*) during the choice test only does not block deprivation-induced preference increase for 3% agarose with 2mM aa incentive in mated females. Experiment conducted at the restrictive temperature of 31°C. Groups: no *Gal4* driver no deprivation (n=48), no driver protein deprived (n=48); *PAM-Gal4* no deprivation (n=48); *PAM-Gal4* protein deprived (n=48). **C** Acute silencing of *TH-Gal4* dopamine neurons blocks deprivation-induced preference increase for 3% agarose with 2mM aa incentive in mated females. Experiment conducted at 31°C. Groups: no *Gal4* driver no deprivation (n=45), no driver protein deprived (n=46); *TH-Gal4* no deprivation (n=46), *TH-Gal4* protein deprived (n=47). **D** Acute silencing of *TH-Gal4* dopamine neurons also blocks deprivation-induced preference changes in a FLAME variant with amino acid in both options (1.5% agarose with 2mM aa incentive and 50mM sucrose vs .5mM aa with 50mM Sucrose in liquid). Experiment conducted at 31°C with mated females. Groups: no *Gal4* driver no deprivation (n=38), no driver protein deprived (n=37); *TH-Gal4* no deprivation (n=41), *TH-Gal4* protein deprived (n=34).

We next tested the acute requirement of dopamine neurons in the PAM (protocerebral anterior medial) region for aa drive, but no effect of genotype (*F*_(1,188)_=0.95, *p*=0.33) nor interaction between genotype and deprivation (*F*_(1,188)_=0.02, *p*=0.88) was observed, while there was a main effect of deprivation (F_(1,188)_=14.9, *p*<0.001; Fig.4B). In these experiments flies were kept at room temperature and then assayed at 31°C for preference, therefore testing the acute requirement of PAM neurons for aa drive during the choice by virtue of acute neuronal silencing with *UAS-shibire^ts^*. The results therefore suggest that PAM dopaminergic activity is not acutely required to drive flies to work for aa.

There are many dopamine neurons outside of the PAM cluster, and to test the requirement of those we used TH-Gal4 to silence these neurons acutely during the feeding choice (Fig.4C). Unlike for PAM-Gal4, this experiment showed a main effect of genotype (*F*_(1,180)_=8.1, *p*=0.005), deprivation (*F*_(1,180)_=12.4, *p*<0.001) and an interaction between those two (*F*_(1,180)_=4.6, *p*=0.034). Furthermore, deprivation significantly drove control flies to consume more hard aa (*p*<0.001, Sidak’s multiple comparison test), but not experimental *TH>shibire^ts^* flies (*p*=0.54). No interaction was found at 20°C, where *shibire^ts^*is inactive (*F*_(1,188)_=0.071, *p*=0.79) and deprived flies of both the control +> *shibire^ts^* (*p*<0.001) and the experimental *TH>shibire^ts^* genotype (*p*<0.001) strongly increased preference for hard aa with deprivation (Fig. S2A), suggesting that it is indeed silencing of *TH-Gal4* neurons that reduces motivation for hard amino acids. To determine whether altered agarose perception in deprived females caused enhanced agarose avoidance and a perceived lack of motivation at 31°C with *TH-Gal4* neurons silencing, we tested agarose avoidance and found no effect of genotype (*F*_(1,250)_=0.060, *p*=0.81) or assay temperature (*F*_(1,250)_=0.44, *p*=0.51; Fig. S2B), supporting an effect of *TH-Gal4* neurons on motivation, and not perception of the agarose barrier.

In above experiments flies choose between aa embedded in agar, or no aa at all, while both choices contained equal sucrose. We next wanted to determine whether an aa deprivation-induced drive for aa would also motivate the flies to choose hard 2mM aa when the liquid solution also contained some aa, albeit at a lower concentration of 0.5mM. Because of these changed assay conditions we reduced the sucrose concentration in both choices to 50mM, to accentuate the aa in the solution. We also reduced the agarose concentration to 1.5%, since the liquid choice now contained aa as well, and under these conditions, undeprived flies displayed preference for the liquid solution in a good dynamic range (Fig. 4D). A *TH>shibre^ts^*silencing experiment again showed a main effect of genotype (*F*_(1,146)_=5.9, *p*=0.017), deprivation (*F*_(1,146)_=5.0, *p*<0.027) and an interaction between those two (*F*_(1,146)_=7.6, *p*=0.007).

We next asked if TH-Gal4 neuronal activity was required for protein motivation specifically. The same *TH>shibre^ts^* silencing experiment showed a main effect of salt deprivation on hard salt consumption, but no effect of genotype or interaction (Fig. S2C). Similarly, wet starvation was a main effect for hard sucrose consumption (*F*_(1,203)_=4.9, *p*<0.001), but genotype was not (*F*_(1,146)_=2.7, *p*=0.10), nor was there an interaction (*F*_(1,203)_=0.87, *p*=0.35). It is worth noting, however, that the preference for hard sucrose was stronger (*p*<0.001, 0.38 difference in median preference) in *+>shibire^ts^* control flies than in *TH>shibire^ts^*experimental flies (*p*=0.012, 0.12 median difference). Therefore, while TH-Gal4 neurons may not be involved in motivation for salt, there may be an involvement in sucrose motivation that might be uncovered with more specific dopaminergic manipulations.

## Discussion

Assays measuring motivation are important to dissect neural and molecular mechanisms relevant to many neuropsychiatric disorders. Here we show that the FLAME is a versatile conflict choice assay for investigating multiple types of nutritional drive. We show that agarose serves as a barrier to feeding, and that flies will overcome this barrier when there is sufficient external incentive as well as internal drive. But does the FLAME really measure a willingness to exert effort – aka “work” – for nutrients? First, we show that flies do not avoid unsolidified agarose (Fig.1B), suggesting that it is not the taste of it that they avoid. Furthermore, we have shown that feeding bouts on liquid food can last many seconds [Mishra 2021], while on 1% agarose flies consume by taking many individual sips of sub-second duration [Itskov 2014], suggesting that it requires more physical effort to feed on solid food. We also show that the harder the gel, the less flies like to consume from it compared to consuming from liquid food (Fig.1A). Given the lack of a masticatory apparatus (teeth and jaw), flies rely on sucking nutrients out of the food source and that would clearly be easier the more liquified the food.

Using the FLAME, we investigated three nutritional incentives: sucrose, salt, and amino acids. We found that each could induce flies to increase their willingness to overcome agarose when flies had undergone corresponding deprivation of the nutrient (Figures 1-3). While each of these incentives could drive willingness to work, they are different on several levels. Sucrose serves as an energy source to flies [Matilla 2017]; salt, while not caloric, is an essential component of multiple physiological processes and is needed for egg laying [walker 2015]; amino acids provide protein, a building block for many processes, which can also be used for calories when necessary [Chatterjee 2021] but is also essential for egg laying [Lee 2008]. Though they have distinct nutritional functions, gustatory attraction for salt, sugar, and amino acids is largely mediated by sweet gustatory receptor neurons (GRNs) expressing Gr64f [Jaeger 2018, Guillemin 2025]. Specific receptors and receptor complexes enable sensing of these diverse substances by the same neurons [Jiao 2008, Dweck 2022, Aryal 2022, Guillemin 2025, Wang 2026]. Even with this shared attractive population of neurons, the motivational drives we investigated are behaviorally separable, and amino acid deprivation did not motivate flies to overcome agarose for sucrose or for salt (Fig.4 G, H). This, of course, makes sense, physiologically speaking, as the deprivation of a specific nutrient should drive organisms to seek said nutrient specifically and not waste energy on foraging or working for plentiful nutrients. It also suggests that higher order processing is involved in the regulation of these specific drives.

Starvation desensitizes the peripheral perception of bitterness and the degree of peripheral inhibition of sweet neurons by bitter sensing neurons [Chu 2014, Le Due 2016], which leads to reduced avoidance of bitter substances [Mishra 2021]. Similarly, we found that amino acid deprivation reduced the avoidance of bitter caffeine (Fig.4E). Mechanosensory neurons can inhibit sweet neurons at the periphery [Jeong 2016]. However, neither of the three nutrient deprivations changed the avoidance of agarose in our assay except for a mild change after salt deprivation in females (Figs.1-3), suggesting that deprivation does not reduce the activity of the mechanosensory neurons that detect agarose hardness [Jeong 2016, Zhang 2016, Sanchez-Alcaniz 2017]. And likewise, while GRN activity could be potentiated in our deprivation paradigms, thus outcompeting the inhibitory mechanosensory input, the shared reliance on sweet GRNs for attraction to our different incentives makes it unlikely that a change in primary mechanosensation is the main mechanism for nutritional drive, given its behavioral specificity.

Mating can also increase a fly’s willingness to work, but not always across metabolic states. While salt and protein drives were both influenced by mating status in our assay, the exact effect was different. We found that for protein drive, mating and deprivation interacted (Fig 3D), which is consistent with previously published evidence that mated female flies are especially sensitive to protein/yeast deprivation [Ribeiro 2010, Corrales-Carvajal 2016, Uchizono 2017, Munch 2022, Wu 2024]. On the other hand, we found that mating increases salt drive across deprivation status but does not interact with the latter (Fig 2D). The independence of the effects of mating status and salt deprivation has been previously investigated once before [Kim 2024], but most previous work on salt consumption has only investigated the effect of mating or deprivation but not their interaction [walker 2015, McDowell 2026, McDowell 2022, Jaeger 2018]. To our knowledge, no *Drosophila* studies have previously tested the interaction of mating status with two different nutritional manipulations in the same assay. By doing so here, we show that mating status selectively interacts with nutritional state rather than globally increasing egg-related nutrient effort.

Understanding how acute dopamine activity regulates valuation, motivation, and willingness to work is an ongoing question in mammalian systems (Tobler 2005, Wang 2021, Hamid 2016, Grove 2022), but these investigations are limited by the scale of the dopamine system. Flies have a numerically simpler, and genetically more accessible dopamine system, and investigations in flies have previously provided knowledge of dopamine function with exquisite resolution. These include the traditional learning functions of dopamine [Aso 2012, Otto 2020, Fisher 2020], as well as emerging roles for dopamine in modulation of ongoing behavior [Kim 2017, Siju 2021]. We show that the potent increase in willingness to work for protein that comes about after deprivation depends on the acute activity of dopamine neurons labelled by *TH-Gal4*, but not the fly’s canonical reward system labeled by *PAM-Gal4* (Fig 4). Our results provide an additional example of behavioral regulation via short-term activity of dopamine neurons, and point to specific neurons that do, and do not, partake in this regulation. Follow up FLAME studies would therefore be an excellent opportunity to determine how acute dopamine activity (whether it be phasic or tonic, whether it enables plasticity or transiently modifies existing circuits, etc.) enables willingness to work on a highly mechanistic level. In the future, we plan to investigate the particular dopaminergic circuitry required for flies to become willing to work for protein and determine the function of their activity. We also expect it to be possible to investigate the circuitry underlying additional drives, even non-ethological, maladaptive ones brought about by rewards such as drugs of abuse.

---

**Supplemental Figure 1 (relating to Figure 1):**
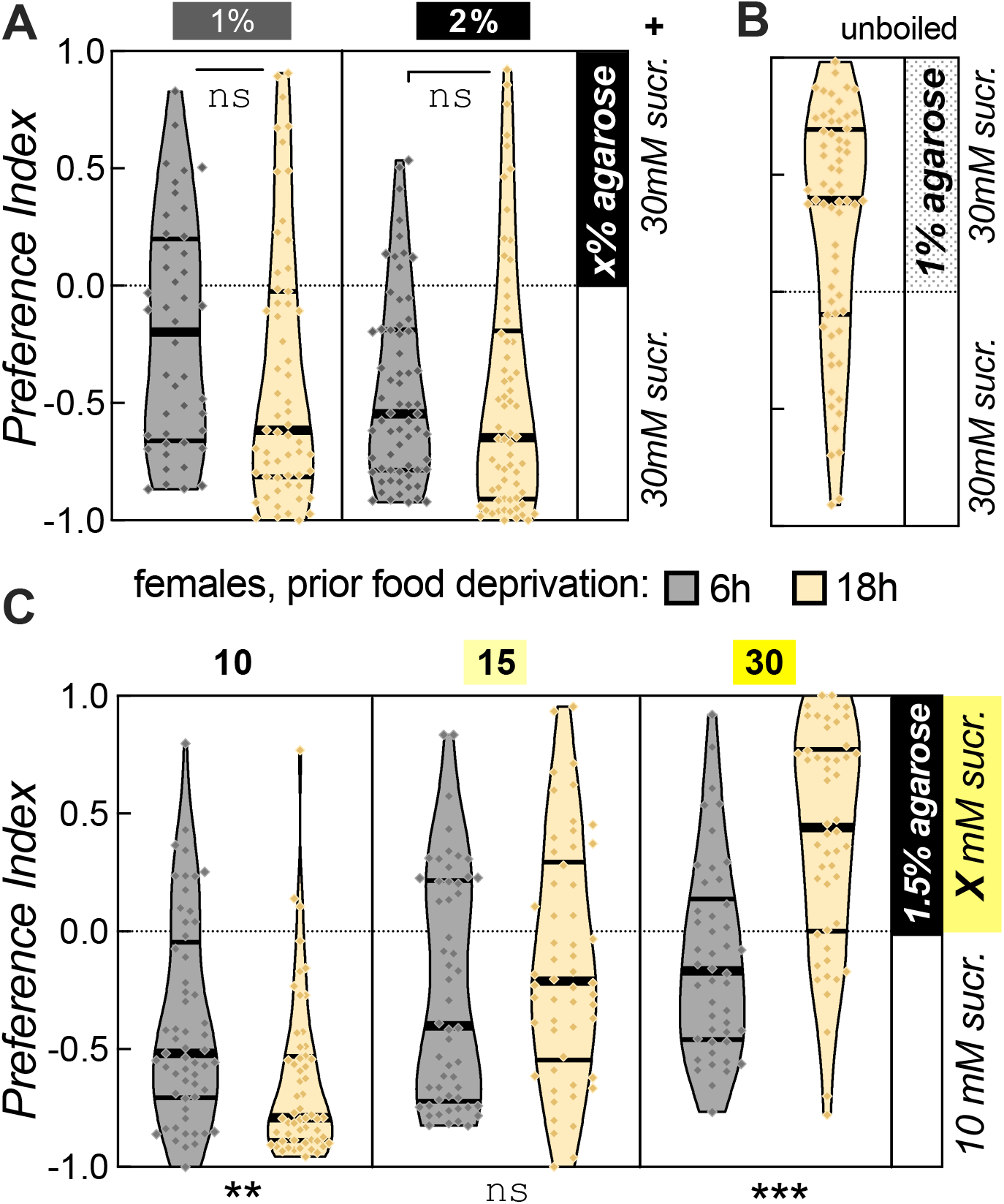
Females exhibit sucrose drive behavior similar to males. **A** Preference for 30mM sucrose in liquid vs in 1% agarose is no different in 18h (yellow) or 6h-food deprived (grey) female wild-type flies (*w* Berlin* for this and subsequent experiments; left). The same is true for 2% agarose (right). Groups: 6h starved 1% agarose (n=44), 18h starved 1% agarose (n=59); 6h starved 2% agarose (n=65), 18h starved 2% agarose (n=66). **B** Non-negative preference for 18h starved female wild-type flies for 30mM sucrose solution containing 1% unboiled agarose vs 30mM sucrose solution without agarose (n=64). **C** Interaction between food deprivation and sucrose incentive in the choice between 10mM liquid sucrose and different sucrose concentrations in hard agar. Only 30mM sucrose drove preference of hard sucrose consumption in 18 h but not 6 h-deprived females. Groups: 6h starved 10mM (n=57), 18h starved 10mM (n=56); 6h starved15mM (n=54), 18h starved 15mM (n=54); 6h starved 30mM (n=46), 18h starved 30mM (n=50).

**Supplemental Figure 2 (relating to Figure 4):**
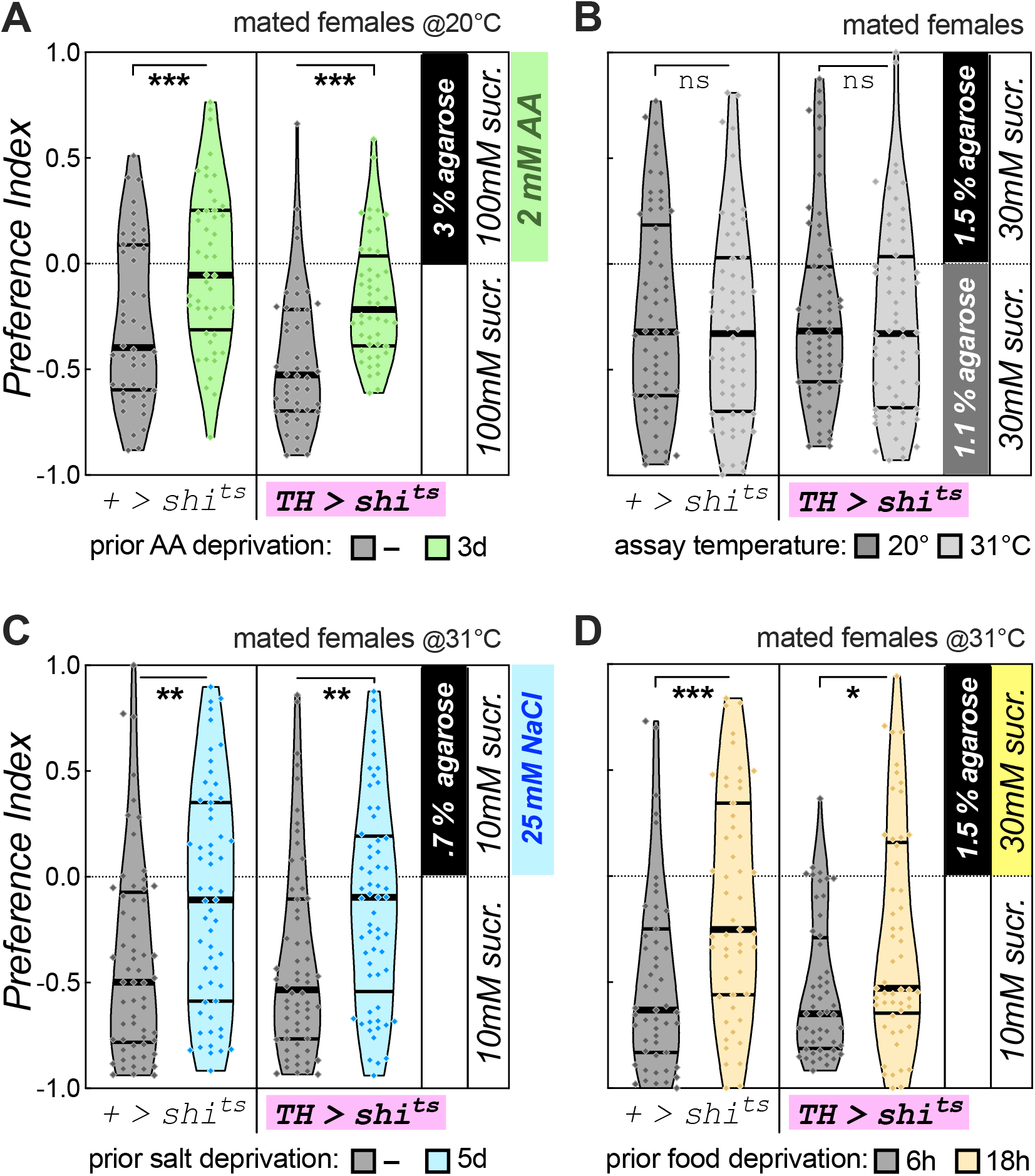
Control experiments for acute *TH-Gal4* neuronal silencing. **A** Preference increase for 3% agarose with 2mM amino acid incentive for mated female flies induced by 3-day protein deprivation is not affected by *TH>shibire^ts^* at the low, ineffective temperature of 20°C. Groups: no driver no deprivation (n=48), no driver protein deprived (n=48); TH-GAL4 no deprivation (n=48), TH-GAL4 protein deprived (n=48). **B** Avoidance of 1.5% agarose vs 1.1% agarose is not affected by temperature, neither in control +>shibirets, nor *TH>shibire^ts^* flies. Groups: no driver 20°C (n=62), no driver 31°C (n=64); *TH-Gal4* 20°C (n=64), *TH-Gal4* 31°C (n=64). **C** Increased preference for 0.7% agarose with 25mM NaCl incentive in mated female flies by a 5-day salt deprivation is not affected by acute silencing of *TH-Gal4* neurons during the choice. Groups: no driver no deprivation (n=61), no driver salt deprived (n=59); *TH-Gal4* no deprivation (n=62), *TH-Gal4* salt deprived (n=61). D Increased preference for hard 30mM Sucrose induced by 18h vs 6h food deprivation is not affected by acute silencing of *TH-Gal4* neurons during the choice (no 2-way ANOVA deprivation x genotype interaction). However, deprivation-induced change is smaller in experimental *TH>shibire^ts^* flies. Groups: no driver 6h starved (n=51), no driver 18h starved (n=51); *TH-Gal4* 6h starved (n=52), *TH-Gal4* 18h starved (n=53).

